# sampleclusteR: A lightweight R package for automated clustering of transcriptomics samples using metadata

**DOI:** 10.1101/2025.04.10.648129

**Authors:** Brandon Coke, Mahesan Niranjan, Rob M. Ewing

## Abstract

**Background:** As technologies for genome-wide gene-expression analysis continue to develop, the databases storing the resulting data have grown accordingly. The Gene Expression Omnibus (GEO) has grown to over 250,000 data series across more than 25,000 omics platforms. Likewise, ArrayExpress is comprised of over 70,000 transcriptome and methylome datasets. Conducting meta-analyses of data from these databases can be challenging, typically requiring extensive manual grouping of samples to identify experimental groups for comparison.

**Results:** Here we present sampleclusteR, a lightweight R package which automates the clustering of gene-expression study samples based on their metadata. To demonstrate the utility of the approach to large scale analysis of GEO data series, 275 GEO data series were analysed using the package. sampleclusteR was able to correctly cluster 4694 of the 5081 samples across the 275 data sets in an unsupervised manner. In addition, 250 datasets from ArrayExpress were analysed by the package with 8547 of the 9154 samples being automatically clustered into correct groups. We show how sampleclusteR can be used to automate analysis of gene-expression datasets by conducting a meta-analysis of multiple GEO data series related to the Wnt signaling pathway. sampleclusteR correctly assigned all samples to the correct experimental groups and identified sets of differentially expressed genes for downstream analysis.

**Conclusions:** sampleclusteR enables large-scale analysis of data from GEO or ArrayExpress by automating the clustering of both GEO and ArrayExpress metadata tables using text mining of their associated metadata.

## INTRODUCTION

The increasing adoption of high-throughput functional genomics technologies has resulted in repositories such as Gene Expression Omnibus (GEO)(Clough et al., 2024) and ArrayExpress (Athar et al., 2019) growing substantially. The wealth of studies now available in these repositories has created rich resources for data-mining, for example integrative analyses and meta-analyses for biomarker discovery (Lazar et al., 2013; Nguyen et al., 2016; Shafi et al., 2019).

A myriad of approaches have been developed to mine and analyse gene-expression datasets in GEO such as ARGEOS (Gavrish et al., 2021) which performs enrichR gene ontology (GO) analysis via GEO2Enrichr (Gundersen et al., 2015) and compendiumdb an R package which streamlines the creation and access of a locally stored compendium of GEO data series (Nandal et al., 2016).

GEOquery (Davis and Meltzer, 2007) is one the most popular R packages for mining and re-analysis of GEO datasets, providing easy-to-use command-line access to GEO Series raw data and metadata. A substantial hurdle, however, when processing large sets of GEO Series, is to automate or semi-automate the clustering of sample groups within the series, such as treatment groups or replicates, so that differentially expressed sets of genes can then be identified. Automating the grouping of GEO Samples through clustering based on their metadata remains a challenge to any large-scale processing of multiple datasets. A previously published R shiny app, GEOracle (Djordjevic et al., 2019), enables users to perform a semi-supervised clustering of GEO data series samples via text-mining the free metadata associated with datasets (Djordjevic et al., 2019). However, several challenges remain for users wanting to process large sets of GEO datasets in an automated manner. Herein we introduce sampleclusteR, an R package for command-line based clustering of samples through analysis of sample annotation and meta-data

### TOOL DESCRIPTION

To address the challenges encountered when processing GEO Series for differential analysis, we developed sampleclusteR with the following features: (1) a command-line interface to facilitate processing of large numbers of GEO Series, (2) reduced disk space requirements since, unlike GEOracle, a cached GEO SQLite database is not stored, (3) fewer R package dependencies compared to GEOracle (19 in GEOracle, 5 in sampleclusteR), (4) the capability to cluster datasets using user-specified metadata (i.e. datasets not in GEO) and data in alternative formats such as Sample and Data Relationship Format (.sdrf) (Dai et al., 2021). This latter feature allows use of formatted metadata from databases such as ArrayExpress and PRIDE(Vizcaíno et al., 2016), thereby extending the utility of sampleclusteR to proteomics datasets. For user convenience we also integrated widely used differential gene-expression analysis tools, limma (Ritchie et al., 2015) and RankProd (Del Carratore et al., 2017), into sampleclusteR to facilitate production of differentially expressed gene (DEG) lists.

The workflow underlying one of the key functions of sampleclusteR is outlined in Figure 1. This function, *geo*.*cluster()*, takes a GEO Series ID as argument and then obtains the metadata and expression data using the GEOquery package (Davis and Meltzer, 2007). It then attempts to cluster the samples using the associated metadata as outlined in Figure 1 via the R packages *rockchalk* and *cluster* (Johnson, 2022; Maechler et al., 2012). If suitable groups of samples are identified, *geo*.*cluster()* outputs a data frame with the samples clustered into groups. If no clusters are identified, the function exits and prompts the user to manually cluster the samples using the associated metadata. In addition to clustering metadata from GEO, sampleclusteR can cluster meta data provided by the user via the *cluster*.*metadata*.*frame()* function which requires a dataframe with the metadata as an input.

**Figure 1.**
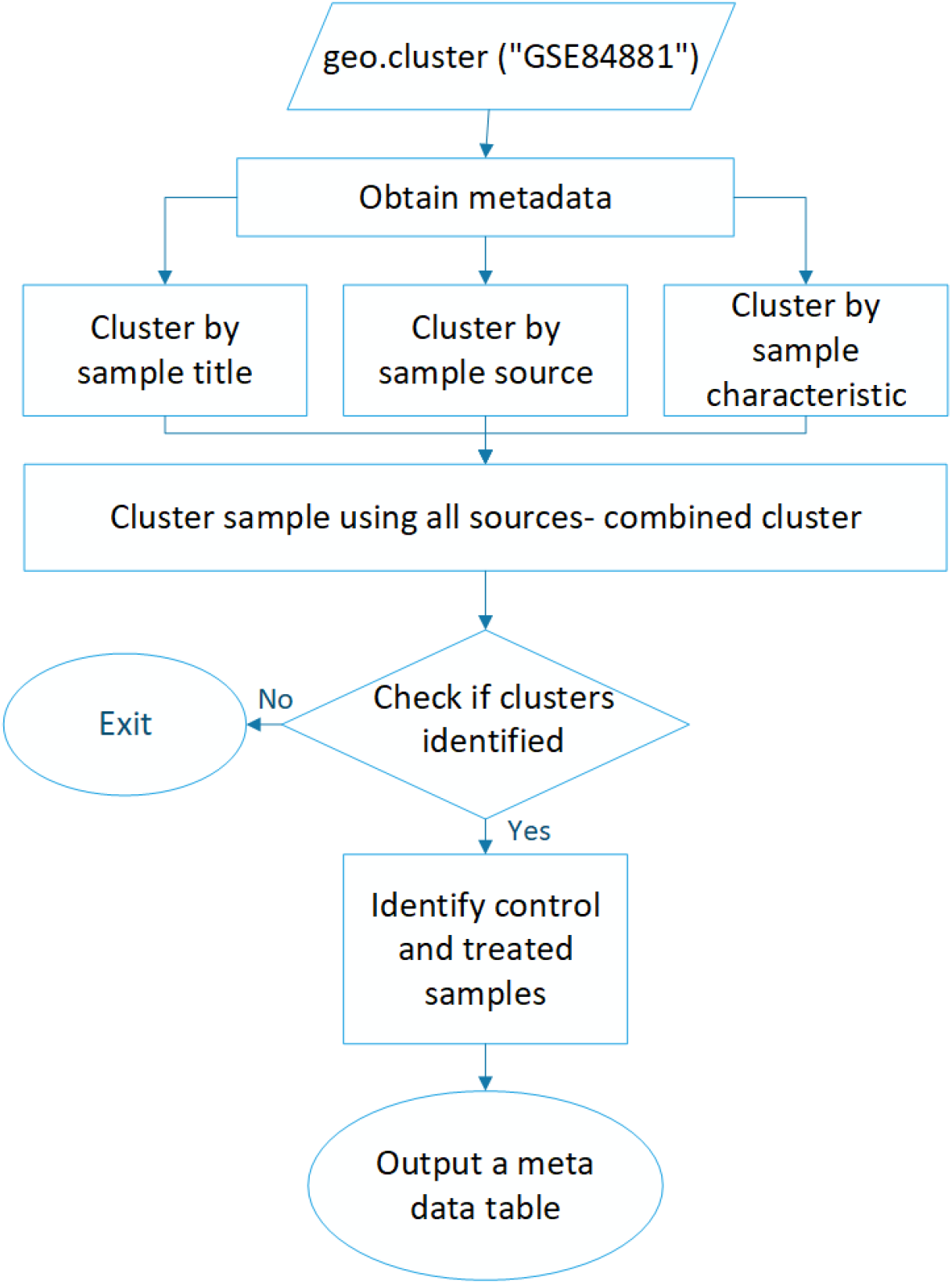
Overview of the *geo.cluster()* function. Metadata associated with a selected GEO data series is obtained via GEOquery. The function then attempts to cluster the samples using the title, source and characteristics assigned to them. If clusters of replicates are identified then a meta data table with predicted sample clusters is produced.

## RESULTS

To demonstrate how sampleclusteR can be used to automate the clustering of a large set of GEO datasets, the *geo*.*cluster()* function was applied to 275 data series across five different gene-expression platforms (Table 1).

**Table 1.** Breakdown of GEO Data Series (studies) used in the evaluation of sampleclusteR.

| Platform ID | Platform | Number of studies |
| --- | --- | --- |
| GPL570 | Affymetrix Human Genome U133 Plus 2.0 Array | 75 |
| GPL10558 | Illumina HumanHT-12 V4.0 expression beadchip | 70 |
| GPL1261 | Mouse Affymetrix Mouse Genome 430 2.0 Array | 40 |
| GPL6480 | Agilent-014850 Whole Human Genome Microarray 4x44K G4112F | 45 |
| GPL13222 | Illumina HiSeq 2000 (Arabidopsis thaliana) | 40 |

In an entirely unsupervised manner, sampleclusteR was able to correctly identify the correct controls and treated samples in 263 of the 275 studies. Out of the 5081 samples from the 275 studies, the function was able to correctly cluster 4694 samples resulting in a sensitivity of 0.924 when using all three features in the metadata as shown in Figure 2. In contrast, lower sensitivities were achieved when using a subset of the available components of the metadata.

**Figure 2.**
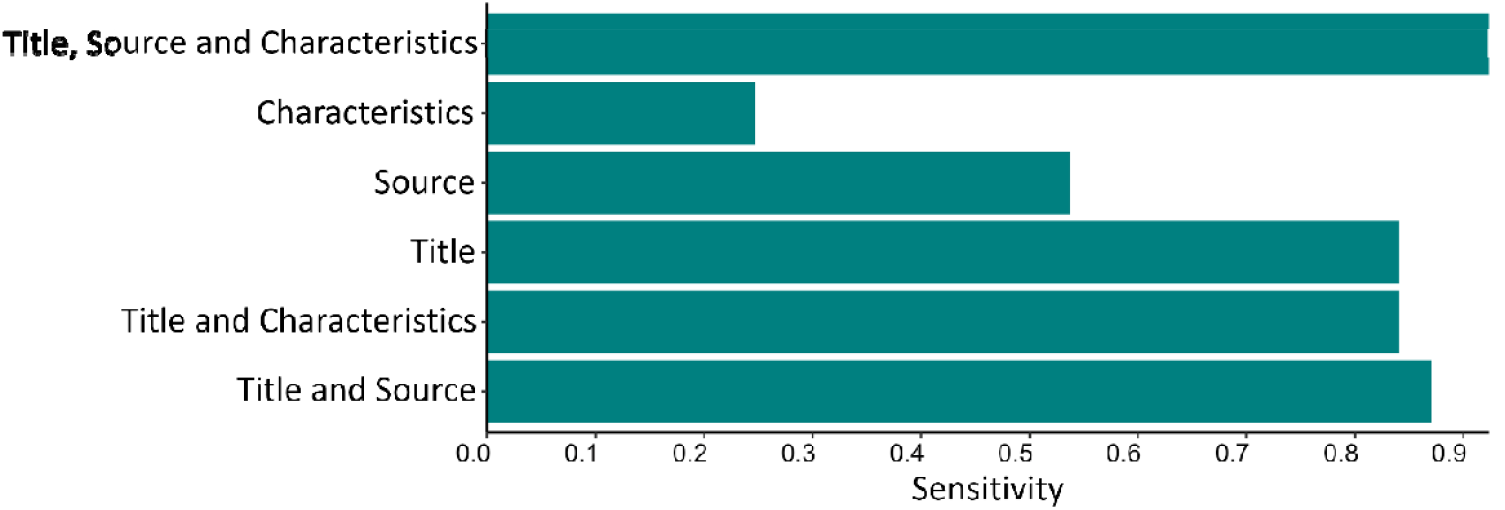
Performance of sample clustering using different sample metadata features. The *geo*.*cluster()* function was used to cluster samples from 275 studies from 5 separate gene-expression platforms in GEO. The ability of the function to correctly cluster the different sample groups was assessed by calculating the sensitivity of the clustering according to the metadata feature (sample title, source or characteristics) combinations used.

One feature which distinguishes sampleclusteR from GEOracle (Djordjevic et al., 2017) is its ability to cluster samples using Sample and Data Relationship Format (.sdrf) formatted metadata used in databases such as ArrayExpress and PRIDE. To assess the ability of the function to automate the clustering of sdrf formatted metadata. 250 studies from ArrayExpress across 5 organisms were clustered using the *sdrf*.*cluster()* function with their numbers outlined in Table 2. Although gene-expression datasets are shared between ArrayExpress and GEO, we selected a set of ArrayExpress studies that are not present in the GEO analysis performed above. In total 9154 samples were included in the analysis. Across the 9154 samples; *sdrf*.*cluster()* was able to correctly cluster 8547 of them (sensitivity of 0.933) with 225 studies out of the 250 being clustered correctly. Samples that were unable to be correctly clustered were primarily due to misspellings of sample descriptions in submitted metadata resulting in misclassifications. Therefore, *sdrf*.*cluster()* can efficiently automate the clustering of samples from ArrayExpress based on their .sdrf formatted metadata table.

**Table 2.**
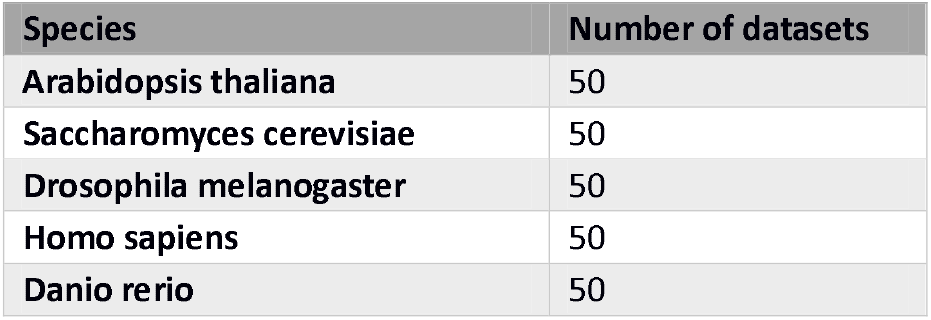
Breakdown of ArrayExpress studies used in assessment.

Finally, we demonstrate how the sampleclusteR package can be used for automated meta-analysis of gene-expression datasets. We show how the *automated*.*analysis()* function can be used to automate both the clustering and analysis of differentially expressed genes. We selected several related studies in which β-catenin, a key protein of the Wnt signalling pathway, has been perturbed over a range of cell types (studies listed in Table 3). These GEO data series were then analysed using the *automated*.*analysis()* function, and we observed that all samples in each of the GEO Series were correctly clustered following automated identification of all treated and control samples. This analysis produced four DEGs lists which then used to produce the Venn diagram and heat map shown in Figure 3. We noted that several canonical markers of Wnt pathway perturbation were identified and indicate these in Figure 3B.

**Table 3.** GEO β-catenin perturbation studies selected for meta-analysis.

| GEO data series | Ectopic or downregulation of B-catenin | Cell line (tissue type) |
| --- | --- | --- |
| GSE17385 | siRNA knockdown | MM1.S (bone marrow) |
| GSE57728 | siRNA knockdown | AsPC1 (Pancreatic) |
| GSE56896 | Ectopic expression | LS174T (Colorectal) |
| GSE44097 | siRNA knockdown | SW480 (Colorectal) |

**Figure 3.**
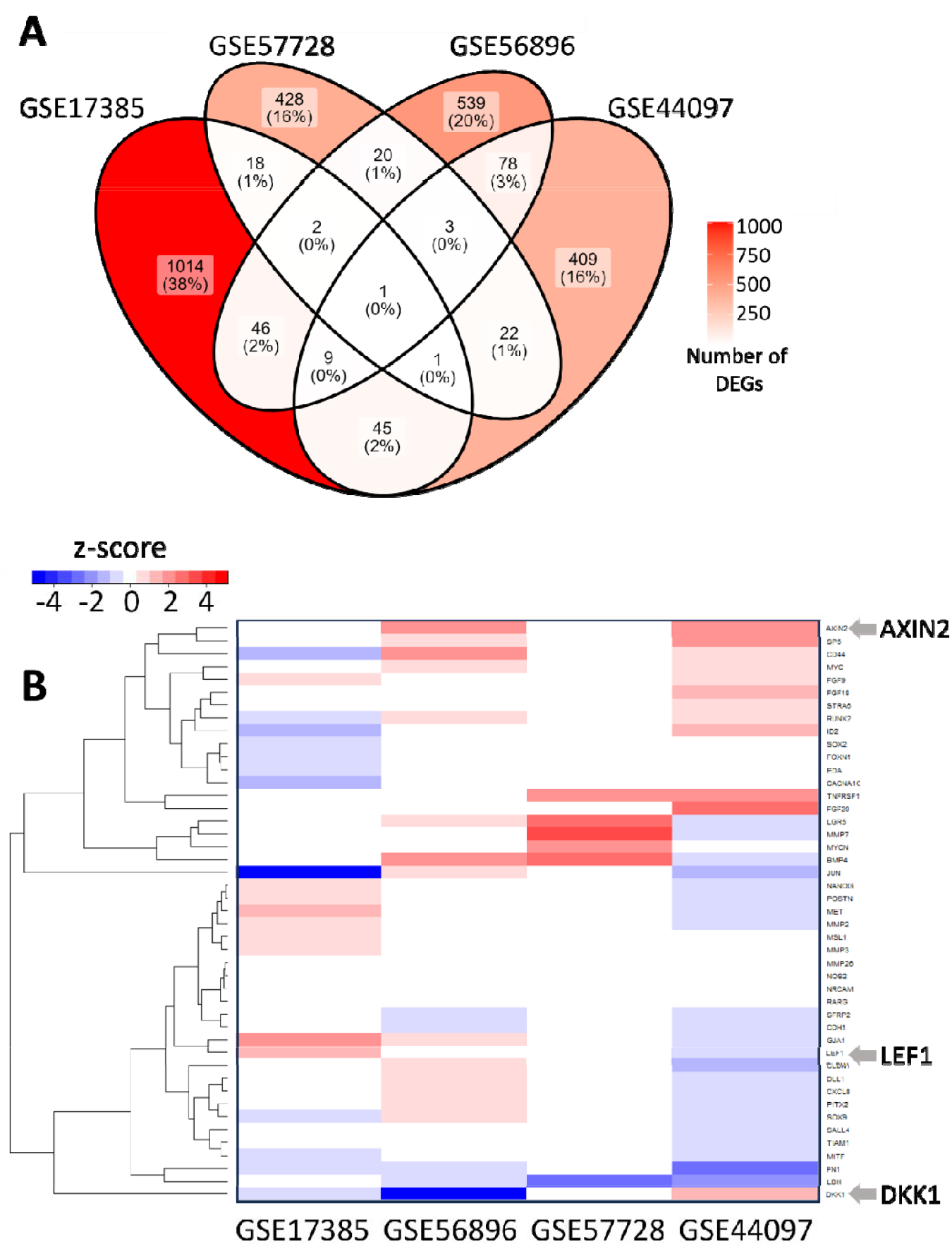
**Using sampleclusteR to perform meta-analysis of multiple GEO data series** sampleclusteR was used to obtain, cluster and then analyse four GEO data series in which expression of a key Wnt pathway regulator (β-catenin) was altered. (A) Venn diagram showing the overlap of DEGs between studies (B) Heatmap of downstream Wnt targets log fold changes (Wnt targets defined by Nusse, 2023), with canonical Wnt markers indicated

### TOOL AVAILABILITY

The git repository (https://github.com/brandoncoke/sampleclusteR) contains all code used in this analysis and installation instructions. The sampleclusteR package can be installed using R via devtools. A Dockerfile is also available which can be used to build a compatible containerised environment. Instructions to run a containerised instance of the package and obtain DEG lists are available on the git repository along with the accompanying Dockerfile. In addition to the Dockerfile, a docker image with a functional instance of the package is also available using R version 4.3.0 along with compatible dependencies. The script used to perform the differential expression analysis (example.analysis.R) is also available from the github site.

## CONCLUSIONS

We developed sampleclusteR to automate the clustering and meta-analysis of gene-expression datasets such as from GEO. Our approach significantly reduces the code required to perform large scale meta-analyses and integrative analyses. Unlike existing tools, sampleclusteR requires fewer dependencies, has a docker container available with a working version of the package, and enables users to analyse their datasets automatically using *limma* or RankProd or alternatively cluster the samples for their own workflow. Finally, the command line interface makes the package amenable for large-scale applications.

## ACKNOWLEDGEMENTS

RME acknowledges Medical Research Council (MR/S01411X/1) and European Commission FP7 (“Oncoprotnet”) for funding. BC and RME acknowledge BBSRC SocoBio DTP for funding.

